# Mechanisms preventing Break-Induced Replication during repair of two-ended DNA double-strand breaks

**DOI:** 10.1101/2020.02.28.969154

**Authors:** Nhung Pham, Zhenxin Yan, Anna Malkova, James E. Haber, Grzegorz Ira

## Abstract

DNA synthesis during homologous recombination (HR) is highly mutagenic and prone to template switches. Two-ended DNA double strand breaks (DSBs) are usually repaired by gene conversion with a short patch of DNA synthesis, thus limiting the mutation load to the vicinity of the DSB. Single-ended DSBs are repaired by Break-Induced Replication (BIR) that involve extensive and mutagenic DNA synthesis spanning even hundreds of kilobases. It remains unknown how mutagenic BIR is suppressed at two-ended DSBs. Here we demonstrate that BIR is suppressed at two-ended DSBs by several proteins coordinating the usage of both DSB ends: ssDNA annealing protein Rad52 and Rad59, D-loop unwinding helicase Mph1, and DSB ends tethering Mre11-Rad50-Xrs2 complex. Finally, BIR is also suppressed when a normally heterochromatic repair template is silenced by Sir2. These findings suggest several mechanisms restricting mutagenic BIR during repair of two-ended DSBs.

## Introduction

Two-ended double-strand breaks (DSBs) are repaired by either nonhomologous end joining (NHEJ) or homologous recombination (HR). In most basic HR pathway by gene conversion, one of the DSB ends invades the homologous template, forming a displacement loop (D-loop) and primes short-patch new DNA synthesis. Repair is completed when the newly synthesized strand is unwound and anneals to the second end of a DSB (**Fig. 1A**). This mechanism called Synthesis-Dependent Strand Annealing (SDSA) is used in mitotic growing cells. Alternatively, the second end of a DSB anneals to the extended D-loop leading to the formation of a double Holliday junction (dHJ), an intermediate common in meiotic recombination and essential for crossover outcomes (Symington, Rothstein et al., 2014). In repair of one-ended DSBs or when homology within the genome is present only for one end of the DSB, this one end invades the template and initiates extensive repair-specific DNA synthesis that can reach even the end of the chromosome. This BIR pathway proceeds via D-loop migration and leads to conservative inheritance of the newly synthesized strands (**Fig. 1A**) (Donnianni & Symington, 2013, Saini, Ramakrishnan et al., 2013). Both SDSA and BIR lead to large increase of point mutations along the entire length of new DNA synthesis (Deem, Keszthelyi et al., 2011, Hicks, Kim et al., 2010, McGill, Shafer et al., 1989, Ponder, Fonville et al., 2005, Saini et al., 2013, Sakofsky, Roberts et al., 2014, Shee, Gibson et al., 2012); however mutation load is far greater in BIR because of the increased length of repair-specific and low fidelity DNA synthesis. In addition, frequent template switches are common in BIR within first 10 kb of strand invasion site (Anand, Tsaponina et al., 2014, Smith, Llorente et al., 2007, Stafa, Donnianni et al., 2014). The increase of point mutations during DSB repair is attributed to the exposure of long single-strand DNA (ssDNA), which is prone to mutagenesis and to the unwinding of newly synthesized strand from its template, which decreases mismatch repair efficiency (Saini et al., 2013, Sakofsky et al., 2014). The frequent template switches in BIR are likely due to the intrinsic instability of the D-loop intermediate (Piazza, Shah et al., 2019, Smith et al., 2007). In diploids, extensive usage of BIR poses a threat to genome integrity by the creation of new mutations, template switches and the loss of heterozygosity. In both diploids and haploids, BIR also generates nonreciprocal translocations by template switching between dispersed homologous repeats (Anand et al., 2014). Indeed, BIR and the related Microhomology-Mediated BIR (MMBIR) are likely the mechanism of many genome instabilities in mammalian cells (Carvalho, Pehlivan et al., 2013, Hastings, Ira et al., 2009, Lee, Carvalho et al., 2007, Li, Roberts et al., 2020, Sakofsky, Ayyar et al., 2015). At least some of the nonrecurrent structural variations that involve large copy number gains are accompanied by hypermutations, implicating a BIR mechanism (Beck, Carvalho et al., 2019).

**Figure 1.**
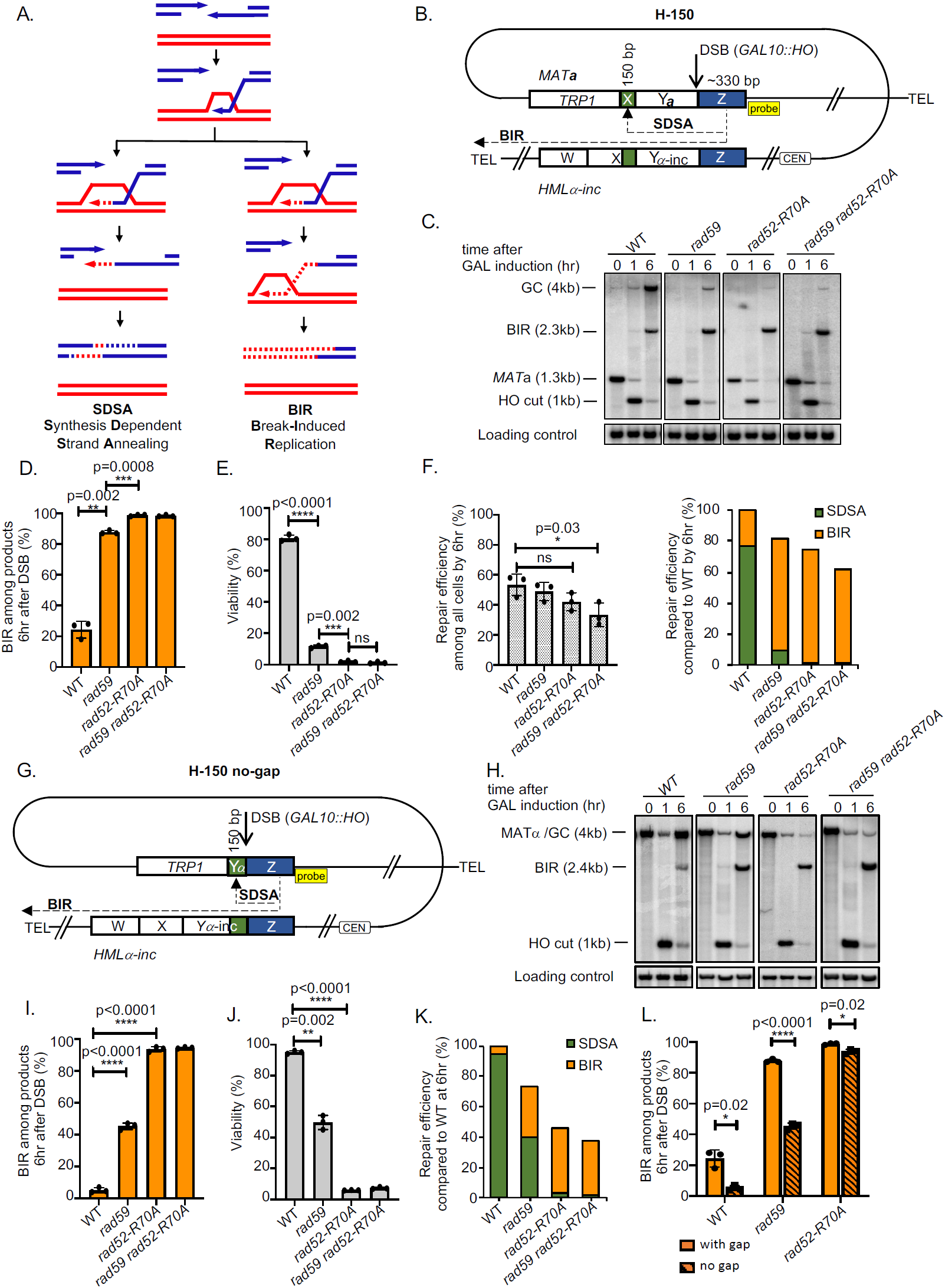
Role of Rad59 and DNA binding domain of Rad52 in suppressing BIR at two-ended DSBs. (A) Models of DSB repair by SDSA and BIR. (B) Schematic of the H-150 assay. Blue boxes depict homologous sequence on one end of the DSB and green boxes depict homology on the other end. A DSB is induced at modified *MAT***a** locus (Chr. III). Strand invasion occurs within the “Z” sequence (blue box) and after copying Yα-inc (dashed line) and shortened X sequence (150 bp), the X sequence is used to capture the second end during SDSA. When copying continues to the end of chromosome *via* BIR (dashed line), an acentric chromosome forms and an unrepaired chromosome segment remains, leading to cell death. (C) Representative Southern blots showing DSB repair products in WT, *rad59*Δ, *rad52-R70A* and *rad59*Δ *rad52-R70A* cells. DNA was digested with Bsp1286I and probed with a *MAT*-distal sequence (yellow box in panel B). (D) Percentage of BIR products among repair outcomes by 6 hr are shown. (mean ± SD; n = 3). Welch’s unpaired t-test was used to determine the p-value in all panels. (E) Viability of indicated strains is shown (mean ± SD; n = 3). (F) Graphs show repair efficiency among all cells (left) and repair efficiency by either SDSA or BIR compared to WT by 6 hr after break induction (right) (G) Schematic of H-150 no-gap assay, where 150 bp of Yα homology is immediately adjacent to the second end of the break. (H) Representative Southern blots showing DSB repair products in indicated mutants in H-150 no-gap assay. (I) Percentage of BIR product among all repair products by 6 hr are shown. (mean ± SD; n = 3). (J) Viability of indicated strains (mean ± SD; n = 3). (K) Repair efficiency of either SDSA or BIR compared to WT of the indicated mutants by 6 hr after break induction. (L) Comparison of BIR product in H-150 assays with gap and without gap of the indicated mutants.

How cells limit the usage of BIR in repair of two-ended breaks is not well understood. When homology on one of two ends of a DSB is short (46-150 bp), BIR contributes or even outcompetes SDSA (Malkova, Naylor et al., 2005, Mehta, Beach et al., 2017). Also, elimination of Mph1, an enzyme disrupting extended D-loops (Piazza et al., 2019, Prakash, Satory et al., 2009, Stafa et al., 2014, Sun, Nandi et al., 2008), likely stabilizes D-loops, and results in the increased BIR outcomes, at least when the homology is short (Mehta et al., 2017).

Here we hypothesized that enzymes ensuring coordination of the usage of two DSB ends in repair may prevent BIR. Proteins important for annealing, stabilizing D-loop and keeping two ends together were examined in this study. We tested whether the elimination of the annealing activity that is required for the second-end capture to form double Holiday junctions or to complete SDSA, can unleash BIR during repair of two-ended breaks. Annealing of ssDNA in yeast is mediated by Rad52 and Rad59 (Mortensen, Bendixen et al., 1996, Sugiyama, New et al., 1998). Both purified enzymes show annealing activity but only Rad52 can anneal complementary ssDNA in presence of RPA (Wu, Sugiyama et al., 2006). Rad59 interacts with Rad52 and stimulates annealing activity of Rad52 in suboptimal conditions, likely by mitigating the negative impact of Rad51 binding to Rad52 on annealing (Davis & Symington, 2001, Gallagher, Pham et al., 2020, Wu, Kantake et al., 2008). In cells, Rad59 seems to be particularly important for annealing of shorter and not completely homologous sequences (Sugawara, Ira et al., 2000). Purified yeast or human Rad52 mediate second end capture to an extended joint molecule (McIlwraith, Vaisman et al., 2005, McIlwraith & West, 2008, McPherson, Lemmers et al., 2004, Nimonkar, Sica et al., 2009, Sugiyama, Kantake et al., 2006). Analysis of recombination intermediates in meiosis also supports the later role of Rad52-mediated annealing in both crossover and noncrossover pathways (Lao, Oh et al., 2008). Besides the role of annealases in controlling BIR during repair of two-ended DSBs, we also tested possible function of Mph1 that disrupts the D-loop and the Mre11-Rad50-Xrs2 complex (MRX) that is responsible for keeping two ends of a DSB together (Kaye, Melo et al., 2004, Lobachev, Vitriol et al., 2004). Finally, we examined the effect of the chromatin state of the donor template on the competition between BIR and SDSA. Multiple intra- and interchromosomal DNA recombination assays with broad range of homology sizes (150 bp to over 100 kb) were used to test the function of these enzymes in competition between SDSA and BIR.

## Results

### Rad59- and Rad52-mediated ssDNA annealing suppresses BIR during repair of DSB

Rad52 is essential for the repair of two-ended DSBs, with two important functions: loading Rad51 onto ssDNA and promoting the second end capture *via* its DNA annealing activity. To test the role of the ssDNA annealing in the choice of repair pathway between BIR and SDSA at two-ended DSBs, a separation-of-function mutant *rad52-R70A* was introduced. This mutant is severely defective in DNA binding and ssDNA annealing but capable of loading Rad51 properly to initiate strand exchange (Shi, Hallwyl et al., 2009). In addition, *rad59*Δ and *rad59*Δ *rad52-R70A* double mutants were constructed. To confirm the deficiency of ssDNA annealing activity in these mutants, we used a single strand annealing (SSA) assay. In this assay, an HO endonuclease-induced DSB is located between two direct and partial *leu2* gene repeats (Vaze, Pellicioli et al., 2002). Upon resection, two complementary ssDNA repeats are exposed and can anneal to each other. The repair is completed by cleavage of the unpaired resected strands between the repeats. Wild type cells complete repair within 6 h, while - as expected - all the mutants show a significant deficiency in SSA (**Fig. S1**).

Gene conversion via SDSA is completed when the newly synthesized strand is displaced from its template and anneals to the second DSB end (**Fig. 1A**). We hypothesized that decreased annealing activity would prevent completion of gene conversion and allow the 3’ end to extend further, to finish the repair by BIR. To test this hypothesis, we used an assay developed previously to study competition between SDSA and BIR (Mehta et al., 2017). This assay involves intrachromosomal mating type switching. A DSB is induced by HO endonuclease at *MAT***a** locus and repaired by recombination with an *HML*α-inc template; the resulting *MAT*α-inc product is not cleaved by HO endonuclease. The difference between regular *MAT* switching and this recombination assay is that the homology size of the second DSB end is shortened from ∼1400 bp to 150 bp (H-150 assay) (**Fig. 1B**). Successful capture of the short second end leads to gene conversion of *MAT***a** to *MAT*α-inc. In the absence of second-end capture, DNA synthesis via migrating D-loop (BIR) continues to the end of the chromosome and produces an acentric chromosome fragment and unrepaired chromosome fragment carrying the centromere, causing cell death. SDSA and BIR products can be distinguished by different restriction fragment sizes on a Southern blot (**Fig. 1C**). We note that one of two possible crossover products has the same size as BIR on a Southern blot. However crossover pathway does not contribute to repair in this assay because the other diagnostic crossover product is not observed. As previously shown in this H-150 assay in wild type cells, SDSA dominated with ∼82% contribution to DSB repair with the remaining 18% being BIR products. Eliminating Rad59 reduced the contribution of the SDSA pathway to ∼13%, while in *rad52-R70A* or *rad59*Δ *rad52R70A* mutants SDSA was further reduced to ∼1% (**Fig. 1C-D**). This result is consistent with a marked decrease of the mutants’ viability, as BIR is a lethal event (**Fig. 1E**). However, not all SDSA events were channeled to BIR as overall product formation (BIR+SDSA) 6 hr after break induction was decreased in the annealing mutants (**Fig. 1F**). This finding suggests that either BIR is decreased or delayed in annealing mutants or that short 150 bp homology on the second end interferes with BIR. Both of these possibilities are tested below.

In mating type switching assay and its derivative H-150 assay (**Fig. 1B**), the Y**a** sequence is replaced by Yα-inc. Because the Y**a** sequence is nonhomologous with the template, the invading strand has to synthesize across a ∼700-bp Yα-inc gap to reach 150 bp homology with the second end. It is possible that this gap and/or the resulting nonhomologous tail on the resected DSB end interfere with engaging the second end in recombination. To eliminate these constraints, we tested recombination between a truncated *MAT*α and *HML*α-inc, where only 150 bp of Yα remains to the left of the DSB. In this assay, homology is present immediately at both DSB ends (H-150 no-gap assay, **Fig. 1G**). In wild type cells, SDSA dominated with >95% DSB repair, while in annealing-defective *rad59*Δ and *rad52-R70A* mutants, SDSA dropped to ∼55% and 1%, respectively, and BIR increased (**Fig. 1H-J**). The double mutant *rad59*Δ *rad52-R70A* showed similar phenotype as the *rad52-R70A* single mutant. Similar to the H-150 assay, the overall repair efficiency decreased in annealing mutants (**Fig. 1K**). Together, these results confirm that annealing activity suppresses BIR at DSBs with short homology, either with or without a nonhomologous gap between homologous sequences. By comparing results obtained in H-150 assays with or without the gap, we also conclude that the gap and/or the presence of a nonhomologous tail likely interfere with the second-end capture in SDSA and therefore increase the usage of BIR, at least when homology on the second end is short (**Fig. 1L**).

### BIR mildly decreases in annealing defective mutants

To test whether BIR itself is affected by any of these mutations, we constructed a new strain, called H-0, where all homology to *HML*α-inc on the second end of the break was eliminated. **(Fig. S2A)**. As confirmed by Southern blot, BIR is the sole pathway of DSB repair and cells do not survive, as BIR is a lethal event in this system (**Fig. S2B,C**). In annealing-defective *rad59*Δ and *rad52-R70A* mutant strains we observed a decrease of BIR efficiency 6 hr after DSB induction to ∼80% and ∼60%, respectively (**Fig. S2D**). Again, the double mutant *rad59*Δ *rad52*-*R70A* showed a similar phenotype as *rad52*-*R70A* single mutant.

The second BIR assay that we used to test the role of annealing activity of Rad59 and Rad52 involves recombination between a truncated and a complete chromosome III (**Fig. S3A**) (Deem, Barker et al., 2008). Here, one end has an extensive homology with the template while the the centromere-distal end shares only 46 bp homology. ∼80% of the cells use BIR for repair after the DSB end with extensive homology invades the template and copies over 100 kb to the end of chromosome. The remaining cells use the short homologous sequence at the second DSB end to complete repair by SDSA. BIR and SDSA products can be distinguished by the presence of *ADE1* markers and resistance to G418 (*KANMX* marker). Consistent with previous assays, *rad52-R70A* or *rad59*Δ *rad52-R70A* derivatives nearly eliminated SDSA among the products (<1%) while BIR increased to about 90% (**Fig. S3B**). Notably, overall DSB repair by BIR is more efficient in this assay when compared to H-0 as only 6-8% of the Rad52 mutant cells lacking proper DNA binding/strand annealing activity failed to repair the break, resulting in chromosome loss. Thus, with longer homology available to the invading strand, the Rad52 DNA binding domain is nearly dispensable for completion of BIR.

One possible function of DNA binding domain of the Rad52 in BIR when homology is short could be related to the stability of the initial D-loop. Annealases related to Rad52 such as RecT or λ beta were shown to promote three-strand exchange (Hall & Kolodner, 1994, Li, Karakousis et al., 1998), an activity that possibly could extend the D-loop. Alternatively, Rad52 could anneal a 3’ invading strand unwound from its template back to disrupted, but still RPA-bound, D-loop. Interestingly, deletion of the Mph1 helicase capable of disrupting D-loop significantly increased BIR efficiency in the *rad52*-*R70A* strain in the H-0 assay, although not to WT levels **(Fig. S2E-F)**. Thus, annealing activity may counteract Mph1 by stabilizing the D-loop, at least when strand invasion is made within short homologous sequences. Model presenting possible functions of Rad52’s DNA binding domain in BIR are summarized in **Fig. S2G**.

### Rad59- and Rad52-mediated annealing suppresses BIR at DSBs in regular *MAT* switching

The above BIR/SDSA competition assays had only 46-150 bp homology on the second DSB end. To test the role of annealing in a system with longer homology, we decided to investigate regular *MAT* switching assay between *MAT***a** and *HML*α-inc. In this assay the homology on the second end is ∼1400 bp (H-1400 assay) (**Fig. 2A**). Southern blot analysis demonstrates that in wild type cells neither BIR nor crossover contribute to the repair, which is nearly 100% efficient. However, annealing-deficient cells *rad59*Δ and particularly *rad52-R70A* increased BIR among products to ∼30% and 60-70%, respectively (**Fig. 2B-C**). As expected, viability is greatly reduced in *rad52-R70A* due to a switch to lethal repair by BIR and inability to complete SDSA (**Fig. 2D**).

**Figure 2.**
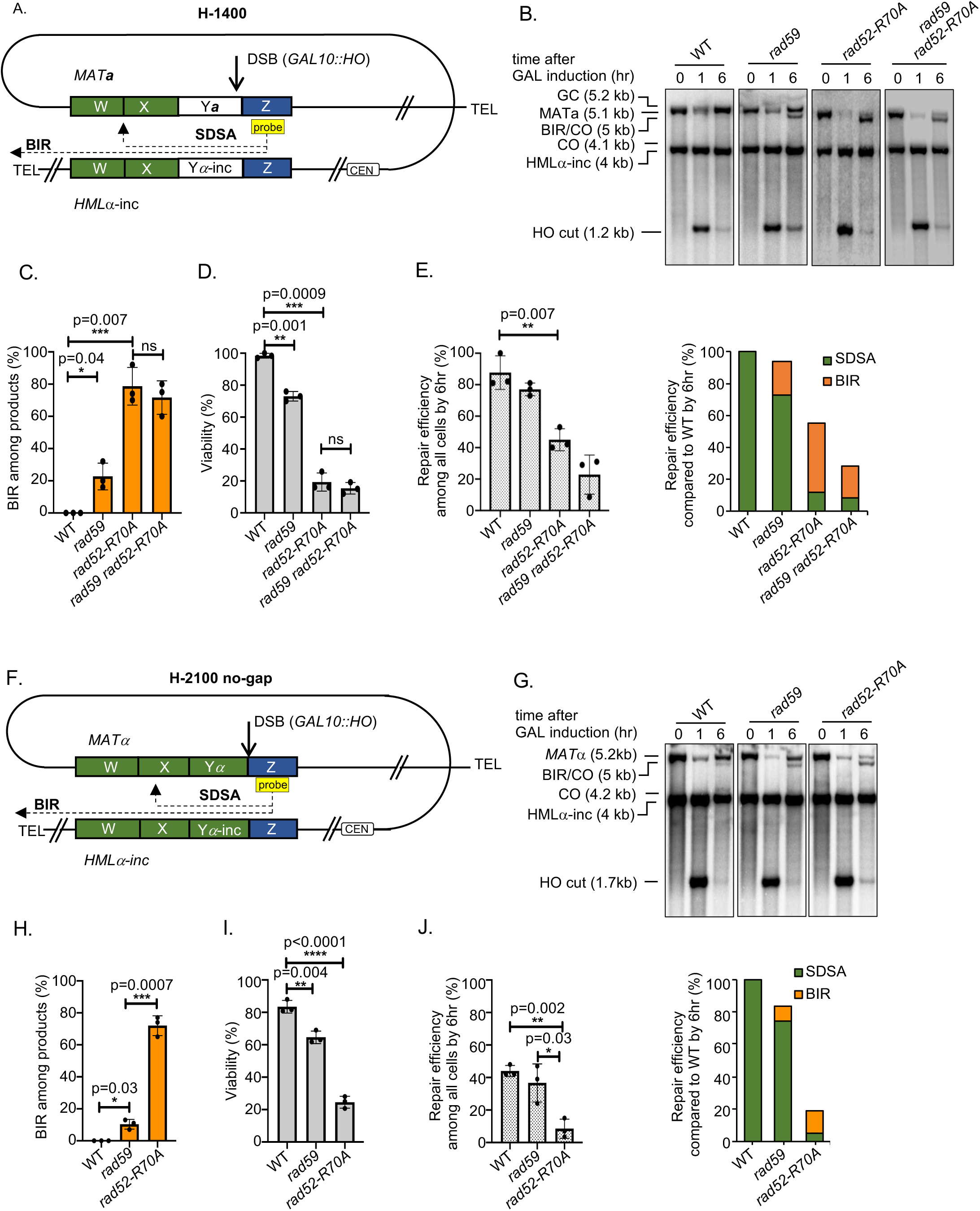
Rad52 and Rad59 suppress BIR during MAT switching. (A) Schematic of the H-1400 assay. A DSB is induced at *MAT***a** and repaired by recombination with *HML*α-inc. There is ∼1400 bp homology between *MAT***a** and *HML*α-inc sequences on the second end (green box). (B) Representative Southern blots showing DSB repair products in WT, *rad59*Δ, *rad52-R70A* and *rad59*Δ *rad52-R70A* cells. DNA was digested with *Xho*I and *Eco*RI and probed with a Z sequence (yellow box). (C) Percentage of BIR product among repair products by 6 hr (mean ± SD; n = 3). Welch’s unpaired t-test was used to determine the p-value in all panels. (D) Viability of indicated strains (mean ± SD; n = 3). (E) Graphs show analysis of repair efficiency among all cells (left) and repair efficiency by either SDSA or BIR compared to WT by 6hr after break induction (right). (F) Schematic of the H-2100 no-gap assay. A DSB is induced at *MAT*α and repaired by recombination with *HML*α-inc. There is ∼2100 bp homology between *MAT*α and *HML*α-inc sequences on the second end (green box). (G) Representative Southern blots showing DSB repair products in WT, *rad59*Δ, and *rad52-R70A*. DNA was digested with *Xho*I and *Eco*RI and probed with a Z sequence (yellow box). (H) Percentage of BIR product among repair products by 6 hr are shown. (mean ± SD; n = 3). (I) Viability of indicated strains (mean ± SD; n = 3). (J) Graphs show repair efficiency among all cells (left) and repair efficiency by either SDSA or BIR compared to WT by 6 hr after break induction (right).

To eliminate the possible constraint of the gap and the nonhomologous Y**a** tail, we tested recombination between *MAT*α and *HML*α-inc where there is no Y sequence heterology (**Fig. 2F**). In this case the homology on the second end is further increased to 2100 bp (H-2100 no-gap) and is present right next to the break. Interestingly, in wild type cells we observed weak bands corresponding to both crossover products (<5%); such crossover products were not observed in recombination between *MAT***a** and *HML*α-inc. Thus, the nonhomologous Y sequence during *MAT* switching prevents lethal intrachromosomal crossovers in ∼5% of the cells. Importantly, in this assay, BIR contribution increases in both *rad59*Δ and in *rad52-R70A* and accordingly viability decreases (**Fig. 2G-I**). We noted above that the BIR product has the same size as the larger of two crossover products. Considering that both crossover products form at the same time and the intensity of the shorter crossover product band is very weak or absent in annealing mutants, we conclude that BIR and not crossover is responsible for the increased intensity of this band in *rad59*Δ and in *rad52-R70A* cells. In this assay long homology is present immediately next to the break on both sides of the break and thus both ends could prime BIR, one toward the telomere and one toward the centromere. However, BIR toward the centromere was not observed in this or any of the other assays tested here. We also note that in H-1400 or H-2100 assays only 15-30% of *rad52-R70A* annealing mutant cells switch to BIR (**Fig. 2E,J**). This level of repair by BIR is far below the one observed in cells that lack any homology on the second end of a DSB (H-0, 60%). Thus, longer homology on the second end, even in the annealing mutants, decreases BIR efficiency. It is possible that with increased homology, both DSB ends might simultaneously invade the template and that this double end invasion would preclude BIR. BIR toward the telomere can still succeed occasionally while BIR toward centromere cannot be completed as previously demonstrated (Morrow, Connelly et al., 1997).

### Increased homology partially alleviates the need of Rad52- and Rad59-mediated ssDNA annealing in recombination and in suppressing BIR

Using intrachromosomal *MAT* switching assays, we found that ssDNA annealing stimulated by Rad59 and Rad52 is important to suppress BIR pathway during repair of two-ended DSBs. All of these assays have homology ranging from 150 bp to ∼2 kb. Next we tested whether Rad52 annealing activity restricts BIR in allelic recombination between two chromosomes III where homology on each end is over 100 kb. To follow different products of DSB repair or chromosome loss in allelic system, chromosome ends were marked with *TRP1, LEU2*, and *KANMX* (**Fig. 3A**). *LEU2* and *KANMX* markers were inserted at subtelomeric regions of the right ends of the two chromosomes. BIR products are viable and can be genetically distinguished from gene conversion with or without crossover and from chromosome loss (**Fig. 3A**). In wild type cells fewer than 1% of cells repair the DSB by BIR. However, BIR increases ∼6 to ∼18 fold in *rad59*Δ, *rad52-R70A* and *rad59*Δ *rad52-R70A* mutant cells (**Fig. 3B**) while viability remains the same as in wild type (100%).

**Figure 3.**
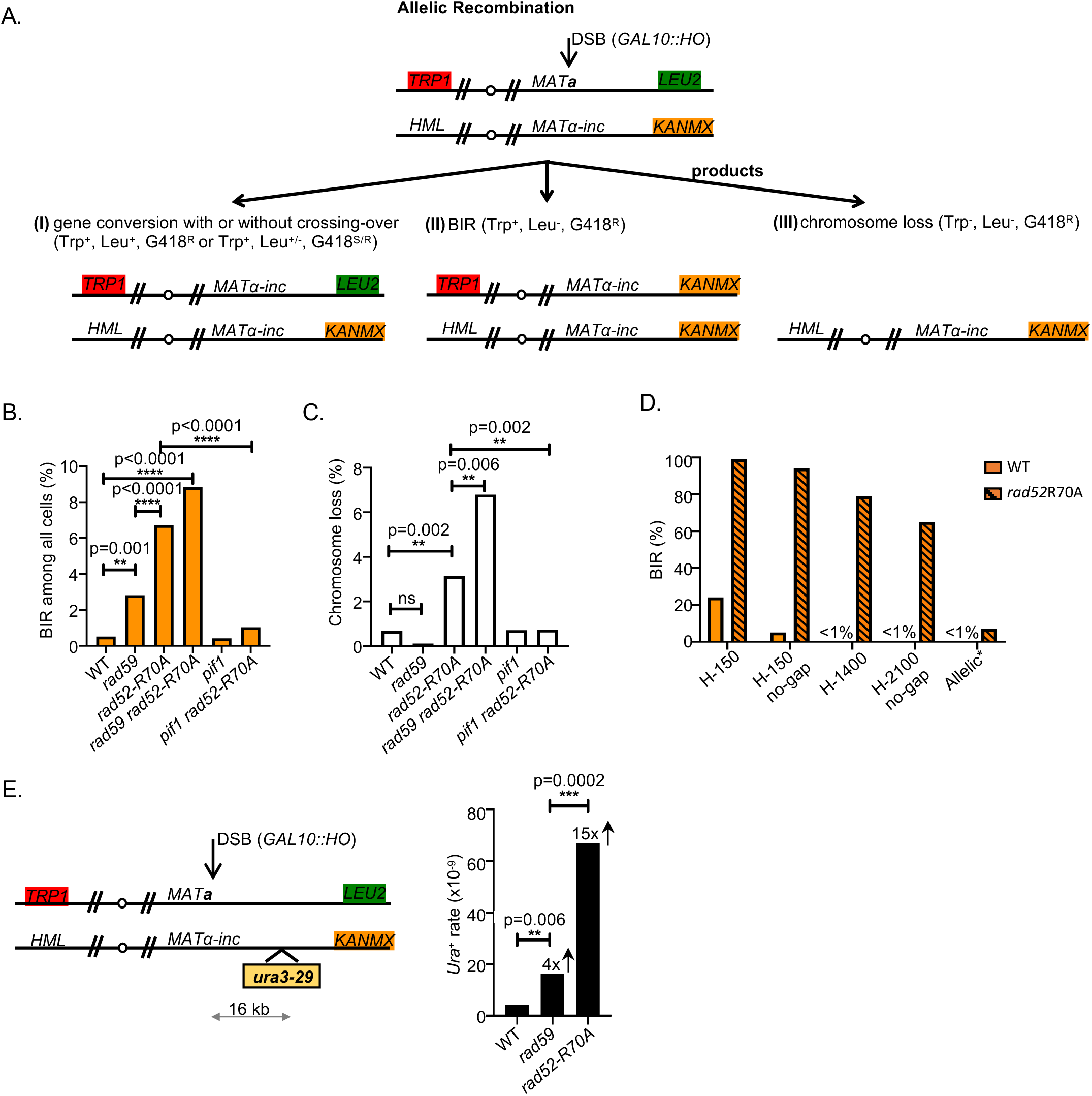
Longer homology mitigates the loss of Rad52 DNA binding domain and Rad59. (A) Schematic of allelic recombination between two chromosomes III. A DSB is induced at *MAT***a** and repaired by recombination with the homologous chromosome carrying *MAT*α-inc. Chromosome ends are marked with *TRP1, LEU2*, and *KANMX* to distinguish different repair products or chromosome loss. (B) Percentage of BIR product among all cells are shown. χ^2^ test is used to determine the p value in this and next panel, number of colonies tested per mutant is indicated in Supplemental Table 1. (C) Percentage of chromosome loss of indicated mutants. (D) Comparison of BIR percentage among the products in different assays in WT and *rad52-R70A* mutant cells. Percentage of repair products are measured at 6 hr after break induction (H-150, H-1400, and H-2100 assays) or scored among plated cells (allelic assay). (E) Schematic of mutation analysis during allelic recombination (left). Mutation reporter cassette is inserted 16 kb away from the break. Mutation rate (right) of WT, *rad59*Δ, *rad52-R70A* was calculated as previously described (Saini et. al., 2013).

The helicase Pif1 is important for BIR as it facilitates D-loop migration (Wilson, Kwon et al., 2013). As expected, elimination of Pif1 greatly reduced BIR in *rad52-R70A* mutant cells during allelic recombination (**Fig. 3B, Table S1**). Compared to *MAT* switching assays, we found that repair in the allelic system was far more efficient in all annealing mutants with only up to 7% chromosome loss (**Fig. 3C**). With all assays taken together, it is clear that increased homology reduces the need of Rad52-mediated annealing for gene conversion (**Fig. 3D**). It is possible that with longer homologous sequences, spontaneous Rad52-independent annealing is sufficient to complete DSB repair (Ozenberger & Roeder, 1991).

### The increased BIR in annealing mutants is accompanied by an increase in mutations

BIR is an extremely mutagenic process owing to its specific mechanism limiting the mismatch repair (Deem et al., 2011, Saini et al., 2013). To test whether BIR contribution to DSB repair between fully homologous chromosomes in annealing mutants is mutagenic, we inserted the *ura3-29* reporter gene 16 kb away from the break on the template chromosome and tested the level of mutations. We observed a 4 and 15-fold increase of mutation rate in *rad59*Δ and *rad52-R70A* mutants respectively when compared to WT cells (**Fig. 3E**). This result is also consistent with the fact that BIR is a highly mutagenic process.

### Role of Mph1 helicase in suppressing BIR at two ended breaks

Mph1 is capable of unwinding D-loops formed during recombination and thus stimulates SDSA pathway in budding and fission yeast (Prakash et al., 2009, Sun et al., 2008). Mph1 is also involved in D-loop unwinding during template switches in BIR (Stafa et al., 2014). Here we constructed mutants defective in both D-loop unwinding and annealing and tested BIR and SDSA contributions to repair of two-ended breaks.

Previously it was demonstrated that elimination of Mph1 increases BIR in repair of two-ended breaks when homology on the second end is short (Mehta et al., 2017), however this increase was much less than observed in annealing mutants. Here we used regular mating type switching where the second end carries 1.4 kb homology (H-1400 assay) to test the role of Mph1 in competition between SDSA and BIR. In wild type cells crossovers are not visible among the products of *MAT* switching (**Fig. 2B**) while in *mph1*Δ two weak crossover bands are observed supporting the role of Mph1 in suppressing crossover outcomes (**Fig. 4A**). The longer crossover band that corresponds also to the BIR product has a slightly stronger intensity than the shorter crossover band suggesting that there is a very mild stimulation of BIR in *mph1*Δ mutant cells. Accordingly, viability of the *mph1*Δ cells is comparable to the wild type cells (**Fig. 4B**). However, elimination of Mph1 in annealing mutants, *rad59*Δ or *rad52-R70A* further increased the BIR contribution to the repair, or nearly entirely switched repair to BIR (**Fig. 4A-D**).

**Figure 4.**
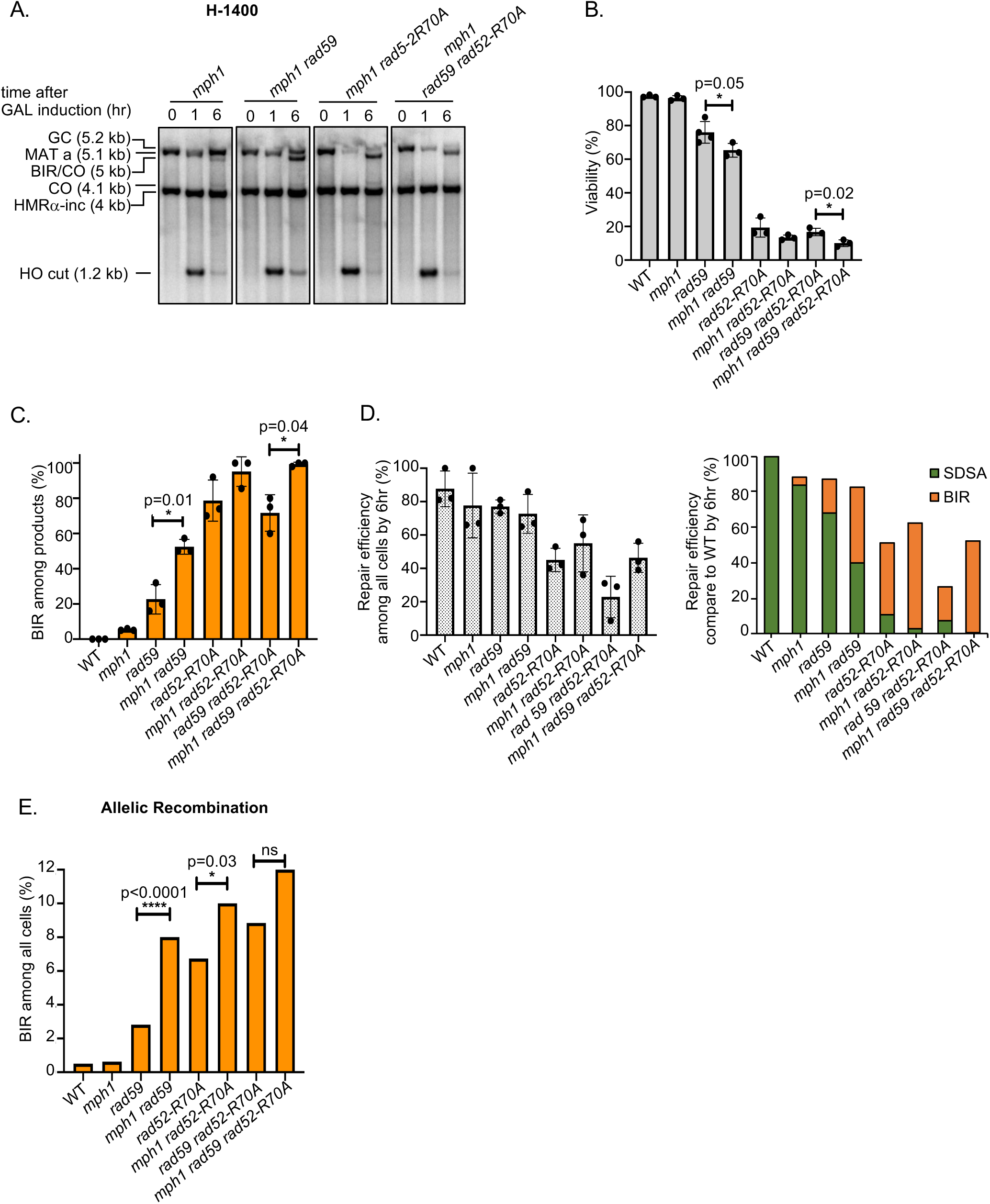
Role of Mph1 in suppressing BIR. Analysis of Mph1’s role in regulating BIR outcomes in mating type switching (A-D) or allelic recombination (E). (A) Representative Southern blots showing DSB repair products of indicated mutants in regular mating type switching (H-1400 assay). (B) Viability of indicated mutants are shown. (mean ± SD; n = 3). Welch’s unpaired t-test was used to determine the p-value in this and next panel. (C) Percentage of BIR product among all repair products by 6 hr are shown. (mean ± SD; n = 3). (D) Graphs show repair efficiency among all cells (left) and repair efficiency by either SDSA or BIR compared to WT by 6 hr after break induction (right). (E) Percentage of BIR among all repair products in allelic recombination. χ^2^ test is used to determine the p value. Number of colonies tested per mutant is indicated in Supplemental Table 1.

In allelic recombination, elimination of Mph1 by itself did not increase BIR events at all; however, loss of Mph1 in annealing mutants increased BIR level to ∼12% in triple *mph1*Δ *rad59*Δ *rad52*-*R70A* mutant cells (**Fig. 4E, Table S1**). We conclude that with extensive homology, Mph1 by itself does not change BIR frequency; however, it suppresses BIR when ssDNA annealing activity is compromised.

### Sir2 suppresses BIR with a heterochromatic donor when homology is short

The *HML* locus serves as template for recombination in *MAT* switching in most of the assays described here (H-150, H-1400, H-2100). *HML* is heterochromatic and silenced by the Sir2 histone deacetylase (reviewed by (Haber, 2012)). Denser chromatin could potentially affect the D-loop migration or D-loop unwinding and therefore impact BIR and SDSA choice. To determine whether the heterochromatic state of the recombination template affects distribution of BIR or SDSA, we constructed *sir2*Δ derivatives of the H-150, H-150 no-gap, and H-1400 assays. The presence of the *HML*α-inc allele prevents cleavage of *HML* when it is unsilenced. In the H-150 assay, the contribution of BIR increased from ∼20% up to ∼55% and viability of *sir2*Δ decreased accordingly, owing to lethality of BIR events (**Fig. 5A**). Similarly, BIR increased by two-fold in the H-150 no-gap system (**Fig. 5B**). In *sir2*Δ cell carrying *HML*α-inc and *MAT***a**, there is a change in mating phenotype. Previous studies have shown that cells expressing both *MAT***a**1 and *HML*α2 genes exhibit some mating-type specific effects on DNA repair (Valencia-Burton, Oki et al., 2006). However, the effect deleting *SIR2* is not related to mating type *per se*, as repair profiles were the same in *sir2*Δ mutant cells that express both *MAT***a**1 and *HML*α2 genes or in the system that expresses only α genes. In regular mating type switching (H-1400 system), we also observe a minimal BIR increase in the *sir2*Δ mutant (∼2%) and no change in viability (**Fig. 5C**). We conclude that closed chromatin structure has a role in suppressing BIR at DSBs, but mostly when homology at the second end is short.

**Figure 5.**
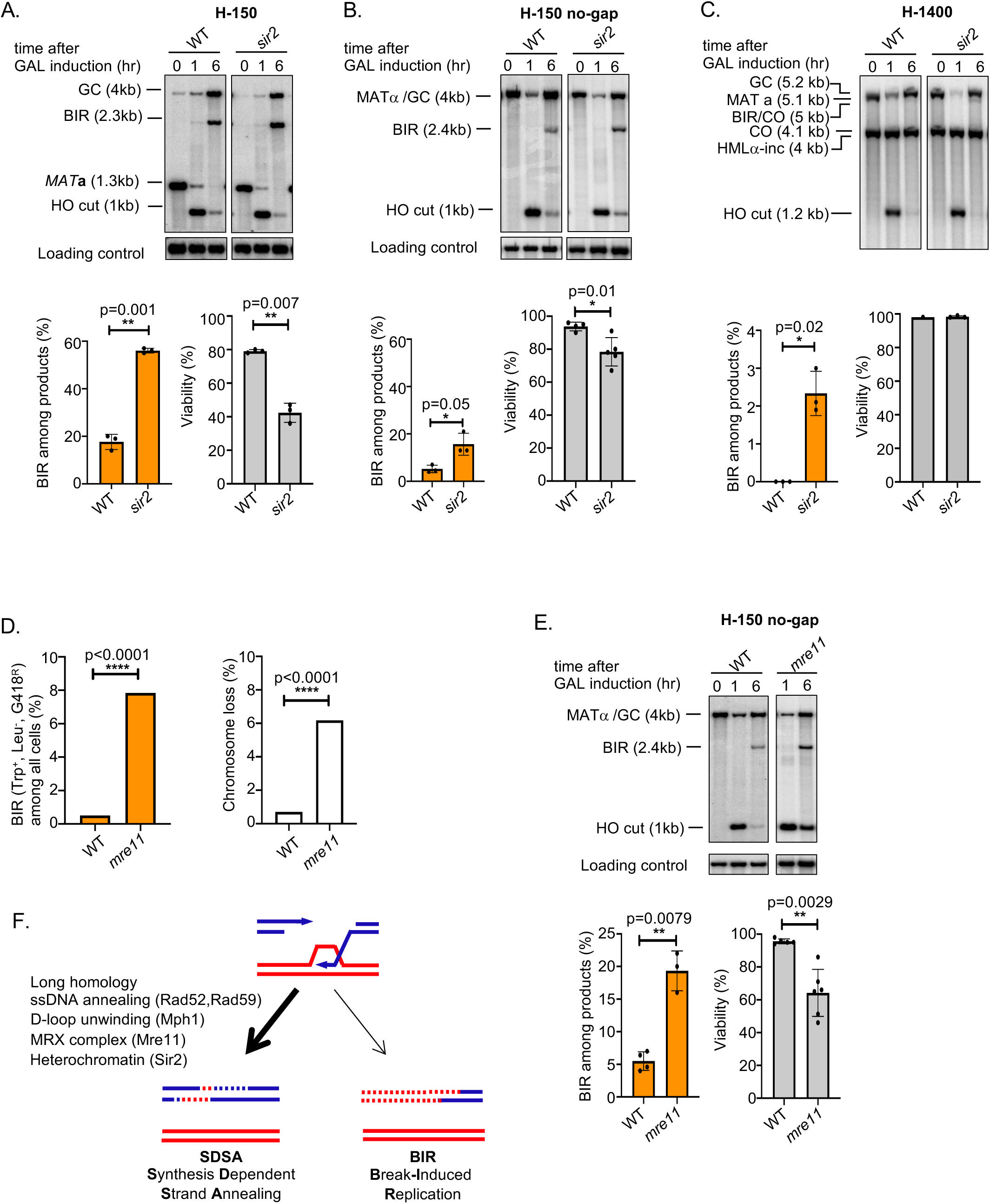
Role of Sir2 and the MRX complex in suppressing BIR during repair of two-ended breaks. (A) Representative Southern blots showing DSB repair products (top), percentage of BIR among repair products (bottom left) and viability (bottom right) of WT and *sir2*Δ in the H-150 assay. (mean ± SD; n = 3). Welch’s unpaired t-test was used to determine the p-value in panels A-C, and E. (B) Representative Southern blots showing DSB repair products (top), percentage of BIR among repair products (bottom left) and viability (bottom right) of WT and *sir2*Δ in the H-150 no-gap assay. (mean ± SD; n ≥ 3). (C) Representative Southern blots showing DSB repair products (top), percentage of BIR among repair products (bottom left) and viability (bottom right) of WT and *sir2*Δ in mating type switching (H-1400) assay (mean ± SD; n = 3). (D) Percentage of BIR and chromosome loss among all cells in WT and *mre11*Δ cells. χ^2^ test is used to determine the p value, number of colonies tested per mutant is indicated in Table S1. (E) Representative Southern blots showing DSB repair products (top), percentage of BIR among repair products (bottom left) and viability (bottom right) of WT and *mre11*Δ in the H-150 no-gap assay. (F) Summary of all factors regulating BIR contribution to repair of two-ended DSBs.

### The MRX complex suppresses BIR during allelic recombination

The MRX complex plays an important role in the initial DSB ends resection (Symington, 2016) and in holding two DSB ends together (Jain, Sugawara et al., 2016a, Kaye et al., 2004, Lobachev et al., 2004). This MRX function may be particularly important in coordination of two DSB ends in repair and therefore in suppressing BIR at two-ended DNA breaks. To test the role of MRX complex we constructed an *mre11*Δ/*mre11*Δ homozygous diploid strain carrying the allelic recombination assay. Full colonies or colonies that showed sectors with a marker pattern typical for BIR products (Leu^-^, Trp^+^, G418^R^) were further evaluated by CHEF and Southern blot with *TRP1* DNA probe to make sure that these are products of expected BIR size and not other genome instability products. Scoring by markers, BIR usage during repair of two-ended DSB increased ∼14-fold (**Fig. 5D)** and 19 of 21 colonies tested by CHEF carried expected BIR products size (**Fig. S4B).** The other two colonies could correspond to BIR-mediated chromosomal rearrangements. The function of MRX complex in suppressing BIR pathway in repair of two-ended DSBs seems to be general as similar phenotype is observed in intrachromosomal H-150 no-gap assay (**Fig. 5E**).

## Discussion

DNA breaks can be repaired by many different pathways of recombination, some of which are mutagenic and normally suppressed. BIR is one of the pathways that drive genome instability as it results in loss of heterozygosity, mutations and nonreciprocal translocations (reviewed by (Sakofsky & Malkova, 2017)). Here we aimed to understand the mechanism that limits the usage of BIR in the repair of two-ended breaks. Several independent mechanisms preventing BIR were identified and are shown in **Figure 5F**. Two-ended breaks are normally repaired by the SDSA mechanism in growing cells, resulting in a short-patch DNA synthesis by polymerase δ (Guo, Hum et al., 2017, Ira, Satory et al., 2006). An essential step in the repair is to engage both ends of a DSB and eliminate the possibility that the invading end primes excessive DNA synthesis up to the end of the chromosome via BIR. We hypothesized that by eliminating activities that coordinate the usage of two ends of a DSB in SDSA, the contribution of BIR would increase. We demonstrate that the loss of ssDNA annealing activity in *rad52-R70A* or *rad59*Δ increases the usage of BIR at two-ended breaks in multiple recombination assays. We propose that compromised ssDNA annealing prevents the second end from annealing to the extended and displaced 3’ strand in the SDSA mechanism or from capturing the extended D-loop in a double-Holiday junction mechanism. In these scenarios a migrating D-loop can reach the end of the chromosome. However decreased annealing has a milder effect on overall repair by gene conversion and on BIR contribution when homology on the second end is extensive. How are two-ended breaks repaired in mutants severely deficient in ssDNA annealing? It is possible that spontaneous annealing is sufficient for gene conversion when homology is long, as proposed earlier in SSA (Ozenberger & Roeder, 1991). Alternatively, with long homology on both sides of the DSB, both ends could invade the template and be resolved as a gene conversion, reducing the chance that D-loop migrate to the end of chromosome. Finally, we cannot exclude the possibility that there is an additional protein capable of weak ssDNA annealing during recombination such as Mcm10 (Mayle, Langston et al., 2019). Notably, in fission yeast Rad52 annealing activity seems to be mostly dispensable for DSB repair as *rad52*Δ cells are resistant to DNA damage so long as cells are capable of loading Rad51 in a Rad52-independent manner (Osman, Dixon et al., 2005).

DNA helicases also coordinate the usage of two ends of a DSB. Mph1 unwinds the extended 3’ strand that subsequently anneals with the second DSB end during SDSA. It is likely that more stable D-loop in the *mph1*Δ mutant cells results in the increased D-loop migration and BIR as previously proposed (Mehta et al., 2017). In addition *mph1*Δ increases BIR when gene conversion is not in competition (Jain, Sugawara et al., 2016b), presumably by preventing unwinding of the initial strand invasion intermediate. BIR suppression by Mph1 in competition with gene conversion is particularly strong with shorter homologous sequences at second DSB end. With longer homology in regular *MAT* switching or allelic recombination, the loss of Mph1 shows increased BIR mostly when ssDNA annealing is compromised (**Fig. 4**). It is likely that besides Mph1, other DNA helicases also coordinate the usage of two DSB ends. Yeast Srs2 is capable of unwinding D-loops, particularly when these are longer (Liu, Ede et al., 2017). In consequence, elimination of Srs2 leads to an increase of crossover products (Ira, Malkova et al., 2003, Mitchel, Lehner et al., 2013, Robert, Dervins et al., 2006). Thus besides Mph1, Srs2 may also suppresses BIR, however we have not tested Srs2 here because it is itself very important for BIR (Elango, Sheng et al., 2017). Lack of the enzymes regulating BIR frequency is partially (e.g. annealing mutants) or entirely (e.g. *mph1*Δ) mitigated by increased homology. This suggests that the control of BIR at two-ended breaks by enzymes coordinating engagement of two DSB ends is very important in recombination between short repeated sequences and thus in organisms with genomes carrying many repetitive elements, such as in humans.

Another enzyme that may coordinate the usage of two ends of a DSB is the MRX complex. Mutants deficient in MRX show frequent (10-20%) separation of two ends of a DSB as monitored by fluorescence microscopy (Jain et al., 2016a, Kaye et al., 2004, Lobachev et al., 2004). This result suggests that MRX is important for holding two DSB ends together. MRX complex also coordinates resection of two DSB ends (Westmoreland, Ma et al., 2009), and synchronous resection of two ends is likely important to engage them in HR at the same time. Indeed, we found that BIR increases over ten-fold in allelic recombination and over 3 folds in H-150 no-gap assay in the absence of Mre11. Also the fact that a gap within template increases BIR contribution (**Fig. 1L**) indicates the importance of keeping two ends together during HR.

*MAT***a** switching normally occurs between the HO-cleaved euchromatic *MAT* locus and a heterochromatic donor, *HML*. We found that Sir2’s maintenance of this heterochromatic state is important for suppression of BIR particularly when homology was short (**Fig. 5A-C**). It is possible that histone deacetylation and highly ordered nucleosomes within *HML*’s chromatin structure interfere with D-loop migration and thus results in easier D-loop disruption. Polδ that is responsible for lagging strand synthesis during replication usually stops DNA synthesis at the point where interaction between DNA and histones is strongest (Smith & Whitehouse, 2012). It is possible that Polδ mediated DNA synthesis in BIR (Donnianni, Zhou et al., 2019) is less effective within dense chromatin. Repressed chromatin structure within repetitive elements and telomeres may prevent BIR during repair of two-ended DSBs.

Altogether this work determines the function of many recombination enzymes in suppressing one of the most mutagenic DSB repair pathways, BIR. Rad52 is highly conserved but in many eukaryotes it plays less prominent role in HR. In these organisms additional annealing enzymes evolved with a role in HR such as FANCA in human, annealing helicase SMARCAL1 in flies, or the BRCA2 homolog in *U. maydis*, Bhr2 (Benitez, Liu et al., 2018, Holsclaw & Sekelsky, 2017, Mazloum & Holloman, 2009). With the exception of Rad59, all the proteins studied here are well conserved between yeast and humans. It is likely that these enzymes prevent mutagenic BIR in a similar way as observed in yeast.

## Materials and Methods

### Strains and plasmids

All stains used in this study are presented in Supplemental Table S2. To study competition between SDSA and BIR using intrachromosomal recombination, we used YAM033 and its derivatives listed in Table S2 (*ho ade3::GAL10::HO HML*α-inc *MAT***a** *hmr::ADE1 bar1::ADE3 nej1::KANMX ade1 leu2,3-112 trp1::hisG ura3-52 thr4 lys5*) (Mehta et al., 2017). To construct strains with different homology size on one end of the DSB at *MAT*, complete/partial deletions of W, X, Yα were made by integrating *Candida glabrata TRP1*, amplified out from YAM033, (primers listed in Table S3). To make the H-150 assay, complete deletion of W and partial deletion of X within *MAT***a**, leaving 150bp, was constructed as described (Mehta et al., 2017). To make the “H-150 no-gap” assay, we created a complete deletion of W, X and most of the Yα sequence within *MAT*α leaving just 150 bp of Yα. For the H-0 assay, all W and X sequences were completely deleted in *MAT***a**. The H-1400 assay represents regular *MAT* switching between *MAT***a** and *HML*α-inc, while the H-2100 assay represents recombination between *MAT*α and *HML*α-inc.

To study the outcomes of the allelic recombination, we used derivatives of JKM139 (*ho hml::ADE1 MAT***a** *hmr::ADE1 ade1 leu2-3,112 lys5 trp1::hisG ura3-52 ade3::GAL10::HO*) and NP633 (*MATα-inc HMLα-inc hmr::ADE1 bar1::ADE3 ade1 leu2,3-112 trp1::hisG ura3-52 thr4 lys5 ade3::GAL10::HO*). To mark the ends of chromosome III in JKM139, a *kiTRP1* marker was inserted at location 17130, and a *LEU2* marker was inserted at position 296670 to create strain NP676, and in NP633 the *KANMX* marker was inserted at position 296670 to generate strain NP643. DNA repair genes listed in Table S2 were deleted in NP676 and NP643 and resulting haploid mutant strains were crossed to obtain diploids for allelic recombination analysis.

To estimate the mutation rate ∼16 kb away from the DSB end during allelic recombination, we first eliminated *ura3-52* from strains NP676 and NP643 using CRISPR/Cas9 methods to construct NP729 and NP728 respectively. Plasmid PL633 was made by cloning gRNA targeting *URA3* (sequence listed in Table S3) into plasmid bRA89 after digestion with *Bpl*I as published (Anand, Beach et al., 2017). Deletion of *ura3-52* was obtained by transforming cells with plasmid PL633 and *URA3* deletion template DNA. Deletion was confirmed by PCR and Sanger sequencing. The *ura3-29-HPH* fragment was generated by PCR amplification using primers listed in Table S3 and inserted ∼16 kb distal to *MAT*α*-inc* in the donor parent strain NP728, resulting in strain NP754. The *ura3-29* mutation reporter (Shcherbakova & Pavlov, 1996) can revert to Ura^+^ via three different base substitutions at one C-G pair. Haploid strains NP729 and NP754 were crossed to obtain diploid NP757 and mutation rates were estimated as previously described (Saini et al., 2013). Diploid strains lacking DNA repair enzymes were constructed for mutation rate analysis as described above and are presented in Table S2.

Mutagenesis of arginine 70 to alanine (R70A) in *RAD52* to make DNA binding-defective *rad52-R70A* mutant was done using CRISPR-Cas9 methods. gRNA targeting *RAD52* was cloned into plasmid bRA90 to construct plasmid PL634. Cells were then transformed with PL634 and *rad52*R70A template DNA to construct the *rad52-R70A* mutant. Mutation was then confirmed by Sanger sequencing.

Single-strand annealing assays were performed in derivatives of YMV45 (*ho hml::ADE1 mata::hisG hmr::ADE1 leu2::leu2(Asp718-SalI)-URA3-pBR332-MATa ade3::GAL10::HO ade1 lys5 ura3-52 trp1::hisG*) (Vaze et al., 2002).

Strains carrying BIR assay between two chromosomes III are derivatives of strain AM1003 (*MAT***a***-LEU2-tel/MAT***a***-inc ade1 met13 ura3 leu2-3,112/leu2 thr4 lys5 hml::ADE1/hml::ADE3 hmr::HYG ade3::GAL-HO FS2::NAT/FS2*) (Deem et. al., 2008).

### Viability assays

To determine the viability in response to a DSB, cells were grown overnight in YEP raffinose medium (1% yeast extract, 2% peptone, 2% raffinose) and ∼100 cells were plated onto YEPD and YEP-galactose plates. Colonies were counted 3-5 days after plating. The proportion of viable cells was estimated by dividing the number of colony-forming units on YEP-galactose plates by that on YEPD plates. Statistical comparison between mutants was performed using Welch’s unpaired t-test and Prism 8 software.

### Growth and induction of DSB in yeast

Yeast cells were grown overnight in YEPD (1% yeast extract, 2% peptone, 2% dextrose) and transferred to YEP raffinose medium (1% yeast extract, 2% peptone, 2% raffinose) for ∼16 hrs. HO was induced when the cell density was ∼1 × 107 cells/ml by adding galactose to a final concentration of 2%. Samples were collected for DNA isolation just prior to and at different time points following addition of galactose.

### Analysis of BIR and SDSA by Southern blot

DNA was isolated by glass bead disruption using a standard phenol extraction protocol. DNA was digested with either *Bsp*1286I (H-0 and H-150, H-150 no-gap system) or *Eco*RI and *Xho*I (H-1400 and H-2100 system) and separated on 0.8% agarose gels. Southern blotting and hybridization with radiolabeled DNA probes was carried out as described previously (Church & Gilbert, 1984). Southern blots were probed with a ^32^P-labeled *MAT***a**-distal or Z fragment. All primers used to amplify the probes are listed in Table S3. All quantitative densitometric analysis was done using ImageQuant TL 5.2 software (GE Healthcare Life Sciences). To control for differences in DNA loaded into each lane, each fragment of interest was normalized to a loading control fragment: *ACT1* fragment (H-0, H-150, and H-150 no-gap) or Z fragment of *HML*α-inc H-1400 and H-2100 system). Percentage of BIR among the products measured at 6 hr after DSB induction was calculated as the percentage of the pixel intensity of the band corresponding to BIR to the total pixel intensity of the bands corresponding to both BIR and SDSA products. In rare cases, a very weak crossover band was observed. In these cases the intensity of shorter crossover band was subtracted from the band that corresponds to both the longer crossover band and BIR. Repair efficiency among all cells of each mutant was calculated as the percentage of normalized pixel intensity of bands corresponding to both BIR and SDSA products at 6hr to the normalized pixel intensity of parental band at 0 hr. Repair efficiency compared to WT was measured as the percentage of normalized pixel intensity of bands corresponding to either BIR or SDSA at 6hr of a mutant to the normalized pixel intensity of bands corresponding to BIR and SDSA at 6 hr of WT cells. Statistical comparison of the percentage of BIR and repair efficiency between different mutants was performed using Welch’s unpaired unpaired t-test (Prism 8 software).

### Analysis of allelic recombination assay

Allelic recombination between chromosomes III was analyzed using diploid strains where each parental haploid strain carried chromosome III with different markers to follow different repair products as described in Figure 3. The assay was performed by growing cells overnight in YEP-raffinose medium and ∼50-70 cells were plated onto YEPD and YEP-galactose plates. After 3-5 days, replica plating on tryptophan dropout, leucine dropout, or plates with YEPD supplemented with G418 to score different repair outcomes. χ^2^ analysis was used to perform statistical comparison between mutants for individual classes of events using Prism 8 software.

### Mutation analysis

To determine mutation frequency associated with BIR, cells were grown overnight in YEPD overnight, followed by growth in YEP-raffinose to the concentration of ∼1 x 107 cells/ml. Breaks were induced by adding galactose to a final concentration of 2%, and cells were incubated for 6 hrs. ∼3 to 5×10^8^ cells were plated on uracil dropout plate(s) before (0 hr) and 6 hr after galactose addition. Number of colonies were counted after 3-5 days. The mutant frequency was calculated as f = m/N (m is the total number of mutants in a culture and N is the total number of cells in a culture. Since the experimental strains did not divide or underwent a single division between 0 h and 6 h, the rate of mutations after galactose treatment was determined using a simplified version of the Drake equation: μ6 = (f6 - f0), where f6 and f0 are the *Ura*^*+*^ mutation frequencies at times 6 h and 0 h, respectively. Rates are reported as the median value and statistical comparisons between median mutation rates were performed using the Mann–Whitney U-test as previously described (Saini et al., 2013).

### Single Strand Annealing analysis

SSA assays between partial *LEU2* gene repeats were previously described (Vaze et al., 2002). DSB induction and viability assays were carried out as described above. For Southern blot analysis, DNA was isolated, digested with *Kpn*I and separated on 0.8% agarose gels. A *LEU2* sequence was used as a probe for SSA repair product and *ACT1* probe was used as loading control.

### CHEF analysis of allelic recombination products product

To analyze BIR repair products scored by plating assay, chromosomal DNA plugs were prepared and separated on 1% agarose gel at 6 V/cm for 48 hrs (initial time = 20s, final time = 30s) by using the CHEF DRII apparatus (Bio-Rad), followed by Southern blotting and hybridization with a *TRP1* sequence as a probe.

### Analysis of allelic BIR

BIR assay using a disomic strain with an extra, truncated copy of chromosome III was previously described (Saini et al., 2013). The BIR assay was performed by growing cells overnight in 2% raffinose medium and ∼100 cells were plated on YEP-galactose plates to induce the break and on YEPD as control. Different repair outcomes were score after 3-5 days by replica plating on adenine dropout, YEPD supplemented with G418 or clonNAT to score different repair outcomes.

## Acknowledgements

We thank members of the Ira laboratory for critical reading of the manuscript. This work was funded by grants from the National Institute of Health (GM080600 and GM125650 to G.I., R35 GM127029 to J.E.H., R35GM127006 to A.M.).

## Author Contributions

N.P. constructed most strains, designed, conducted, and analyzed data. Z.Y. constructed some Rad52 point mutants. N.P., A.M. and J.E.H. and G.I. designed experiments, analyzed the data and wrote the manuscript.

